# Single-cell molecular and cellular architecture of the mouse neurohypophysis

**DOI:** 10.1101/744466

**Authors:** Qiyu Chen, Dena Leshkowitz, Janna Blechman, Gil Levkowitz

## Abstract

The neurohypophysis (NH), located at the posterior lobe of the pituitary, is a major neuroendocrine tissue, which mediates osmotic balance, blood pressure, reproduction, and lactation by means of releasing the neurohormones oxytocin and arginine-vasopressin from the brain into the peripheral blood circulation. The major cellular components of the NH are hypothalamic axonal termini, fenestrated endothelia and pituicytes, the resident astroglia. However, despite the physiological importance of the NH, the exact molecular signature defining neurohypophyseal cell types and in particular the pituicytes, remains unclear. Using single cell RNA sequencing, we captured seven distinct cell types in the NH and intermediate lobe (IL) of adult male mouse. We revealed novel pituicyte markers showing higher specificity than previously reported. Single molecule *in situ* hybridization revealed spatial organization of the major cell types implying intercellular communications. We present a comprehensive molecular and cellular characterization of neurohypophyseal cell-types serving as a valuable resource for further functional research.

**Significance Statement:** The neurohypophysis (NH) is a major neuroendocrine interface, which allows the brain to regulate the function of peripheral organs in response to specific physiological demands. Despite its importance, a comprehensive molecular description of cell identities in the NH is still lacking. Utilizing single cell RNA sequencing technology, we identified the transcriptomes of five major neurohypophyseal cell types in the adult male mice and mapped the spatial distribution of selected cell types *in situ*. We revealed an unexpected cellular heterogeneity of the neurohypophysis and provide novel molecular markers for neurohypophyseal cell types with higher specificity than previously reported.

## Introduction

The pituitary, also dubbed the hypophysis, is the master endocrine gland that is localized at the base of the hypothalamus in all vertebrate species. It is composed of the adenohypophysis (AH) and the neurohypophysis (NH), also known as the anterior and posterior pituitary, respectively. The mammalian pituitary consists an additional anatomically discernable tissue, the intermediate lobe (IL), which is located between the NH and AH. However, the IL is not as distinguished as in the pituitary of human and some non-mammalian vertebrates, including zebrafish (Larkin and Ansorge, 2017; Norris et al., 2013; Wircer et al., 2016). The hypothalamo-neurohypophyseal system (HNS) encompasses hypothalamic magnocellular neurons residing in the paraventricular nucleus (PVN) and supraoptic nucleus (SON) and project their axons in to the neurohypophysis. Thus, two neuropeptides, oxytocin (OXT) and arginine-vasopressin (AVP) are produced in magnocellular neurons, transported along neurohypophyseal-projecting axons and released into the general blood circulation through the neurohypophyseal capillary plexus (Murphy et al., 2012). Circulating OXT and AVP neurohormones affect the physiological function of peripheral organs such as the kidney, mammary gland and the uterus. Specifically, AVP regulates osmotic balance and blood pressure (Donadon et al., 2018; Olazábal, 2018; Wircer et al., 2016), while OXT is mainly known due to its effects on reproduction organs (Lee et al., 2009).

Unlike the AH, which serves as a hormone-secreting gland, the NH is a neural tissue, which serves as a neuroendocrine interface between AVP and OXT axonal projections and the permeable capillary network of fenestrated endothelia (Robinson and Verbalis, 2003). This neuro-vascular interface also contains the pituicytes – specialized neurohypophyseal astroglia, which occupy approximately 50% of the neurohypophyseal total volume (Bucy, 1930; Pow et al., 1989). Pituicytes engulf HNS axonal swellings and their terminal buttons, and are in close contact with the basal laminar and vascular endothelia (Miyata, 2017; Robinson and Verbalis, 2003). Based on their dynamic morphological plasticity during lactation and in response to chronic dehydration, it has been suggested that the pituicytes mediate neurohormones passage through the fenestrated capillaries serving as a physical gateway between the axons and the perivascular space (Hatton, 1988; Wittkowski, 1998). Recently, we reported that during development, pituicyte-derived factors regulate the decision of zebrafish NH vasculature to adopt a permeable endothelial fate instead of forming a BBB (Anbalagan, 2018). The early definition of pituicytes was based on histochemical staining with silver carbonate and hematoxylin & eosin (Bucy, 1930; Liss, 1958; Dellmann and Sikora, 1981). Thus, different sub-types of pituicytes have been defined by their fibrous, ependymal (with cilia or microvilli), oncocytic morphologies or by ultrastructure of organelle contents, such as dark and pale pituicytes due to high/low density contents of cytoplasmic matrix and organelles and granular pituicytes containing numerous cytosegregosome type dense bodies (Anbalagan et al., 2018; Seyama et al., 1980; Wittkowski, 1986). However, there is a very little knowledge of pituicyte-specific genes. Consequently, mammalian pituicytes have been so far labelled with astroglial markers, such as apolipoprotein E (APOE), glia fibrillary acidic protein (GFAP), S100β, vimentin (VIM) and connexin43 (Cx43/GJA1), all of which are general astrocytic markers, which are also expressed in other cell types (Boyles et al., 1985; Cocchia, 2004; Marin et al., 1989; Suess and Pliška, 1981; Yamamoto and Nagy, 1993). Moreover, defining and visualizing pituicytes by co-expression of the above genes is not informative as these markers only partially overlap (Wei et al., 2009). Hence, the exact definition of pituicyte cell-type and/or sub-type remains ambiguous. Finally, other neurohypophyseal cell types might not have been detected in published bulk neurohypophyseal transcriptomic data (Hindmarch et al., 2006).

Recent technological revolution enables high-resolution studies for transcriptome patterns in heterogeneous cell population. Single cell RNA sequencing (scRNA-Seq) allows dissecting cell types that are previously hidden due to identical histology, same genetic marker and adjacent location within a complex tissue (Potter, 2018). This technology enables hundreds and thousands of single cells being processed at once, therefore deliver high-throughput, and highly efficient analysis of cell heterogeneity. In this study, we utilized scRNA-Seq to unravel the cell heterogeneity of the neurohypophysis. Seven major cell types in the NH and IL of adult male mouse were identified. We present a comprehensive view of the molecular landscape as well as spatial organization of NH and IL cell types, hence providing valuable resources for studying their specific cellular and physiological functions.

## Materials and Methods

### Experimental Design

Three month-old male C57/BL6 and *Cx3cr1*-GFP mice (Jung et al., 2000) were used in this study. All experimental procedures were approved by the Weizmann Institute’s Institutional Animal Care and Use Committee (IACUC).

### Single cell dissociation

Two independent groups of five C57/BL6 mice were sacrificed by decapitation and the NH were dissected and collected into ice cold 1 ml of Magnesium- and Calcium-free HBS -/- buffer [20mM HEPES-buffered saline 145mM NaCl, 5.4mM KCl, 20mM Glucose, pH 7.2] (Dieck, 1999). NH tissues were then transferred to ice cold Phosphate Buffered Saline (PBS) containing Magnesium and Calcium [HyClone, GE Healthcare, USA], treated with 50ng/µl Liberase TM [Roche, USA] for 12mins at 37 °C and further dissociated by incubating in HBS-/- buffer containing 0.15mg/ml Papain (Sigma, USA) and 10U/ml DNase I (Invitrogen, USA) for 8mins at 37 °C. The reaction was stopped by adding heat inactivated fetal bovine serum (HI-FBS) [Hyclone, USA] to reach final concentration of 5%. To obtain single cell resuspension, the loosened tissues were collected and passed through a 40µm nylon mesh in 800µl resuspension buffer [Leibovitz L-15 with 0.3mM Glutamine (Gibco, Thermal Fisher, USA), 0.5% of Penicillin Streptomycin solution (Gibco, Thermo Fisher, USA), 1% HI-FBS, 0.04% BSA]. Cell number, survival rate, clarity and singularity were checked by Trypan Blue staining followed by hemocytometer counting.

### Single cell RNA sequencing

Single cell RNA sequencing was performed with 10X Genomics Chromium Single Cell Kit Version 2. Two independent samples, each containing 600-800 cells/ml, which had approximately 70% survival rate and very few debris were used to form droplets containing single cell and barcoded-beads. The targeted recovery was 4000 cells per sample. The subsequent cDNA synthesis and library preparations were conducted according to manufacturer’s protocol [10x Genomics, USA]. Two libraries were then indexed and pooled for sequencing using a NextSeq 500 High Output v2 kit (75 cycles) [Illumina, USA] according to the manufacturer’s instructions. Four lanes were used with R1 26 cycles and R2 58 cycles.

### Data and software availability

The accession number for the neurohypophysis single cell transcriptome reported in this paper is Gene Expression Omnibus (GEO) (www.ncbi.nlm.nih.gov/geo): GSE135704.

### Statistical Analyses

Sequences data was demultiplexed using Illumina bcl2fastq. Each of the samples was analyzed by Cellranger (version 2.0.0), run with the option --force-cells=1500 and using the 10X prebuilt mm10 reference database version 1.2.0. The outputs from CellRanger were further analyzed using the Seurat package V2.3 (Butler et al., 2018) and R 3.5. Using Seurat, we performed gene filtering (gene must appear in 3 cells of a sample) and merging of the cells of both samples to one set. Cell filtering was based on the number of genes per cell (must be between 400 and 5000), the number of UMI counts per cell (between 1000 and 10,000), and the percent of mitochondria genes lower than 0.25 percent. 11 clusters were created with 900 variable genes and 11 PCs. The cluster names were replaced with the cell type identity based on the differentially expressed genes (marker genes).

### Whole mount in situ hybridization (WISH) and immunostaining

3-month-old C57BL6 mice were perfused and fixed by 2% PFA for 10mins and fixed in 4% PFA for 24h at 4 °C in the dark. WISH was performed as described in (Machluf and Levkowitz, 2011) with prolonged proteinase K treatment of 45min. Tissues were post fixed in 4% PFA for 20mins at room temperature and washed 3×15 min PBS-Tx (Triton X100; 0.3%). Subsequent Immunostaining of WISH samples was performed following re-blocking in blocking buffer [10% Lamb serum, 0.3% TritonX-100, 1% DMSO in PBS] for 1h. Primary antibody staining was performed at 4 °C overnight. After 3×30 min PBS-Tx wash, the samples were incubated with 1:200 secondary antibody at 4 °C overnight, followed by 3×30 min PBS-Tx wash and mounting in 75% glycerol. Imaging of WISH samples was performed using Zeiss LSM 800 confocal microscope with oil immersion 40x objective. Whole z-stack maximum intensity projections were generated by Fiji-ImageJ software.

### Cryotomy and smFISH

C57BL6/Cx3cr1-GFP transgenic mice were sacrificed by decapitation. The whole pituitary was quickly dissected and fixed in 1% PFA containing 30% sucrose overnight at 4 °C. The fixed tissue was then washed and equilibrated in half Tissue-Tek O.C.T Compound [Sakura, USA] and half 60% sucrose (final 30%) mixture before positioned inside a plastic mold with only O.C.T compound and frozen by burying in dry ice powder. After the whole block turned opaque, it was stored at −80 °C in a sealed plastic bag in the dark. Prior to cryotomy, the embedded O.C.T block was first equilibrated inside the Cryostat machine [Leica, Germany] to −25 °C for 30mins followed by cryo-sectioning (7µm) and slice collection on 22×22 mm glass coverslips #1 [Thermo Scientific Menzel, USA], precoated with 0.01% L-lysine (Sigma), and stored at −80 °C in a parafilm sealed 6-well plate in the dark for up to a month before further digestion and prehybridization steps. smFISH was conducted as described in (Ji and van Oudenaarden, 2012) with the exception that the formamide concentration was increased to 30% for prehybridization and washing. Tissue sections were mounted on Prolong Gold antifade mountant [Thermo Fisher, USA] and images were captured using a wide-field fluorescent microscope [Nikon Eclipse Ti-E, Japan] with a cooled CCD camera equipped with oil immersion 60x objective.

### Vibratome sections

Pituitary from Cx3cr1-GFP mouse was dissected on ice and fixed in 4% PFA overnight at 4 °C. After washing, the pituitary was embedded in 3% Nobel Agarose [BD Biosciences, USA] on ice. 50µm coronal sections were cut using a Leica VT1000 S vibrating blade microtome [Leica, Germany] and then mounted with Aqua-Poly/Mount [Polysciences, USA]. The sections were then imaged using Zeiss LSM 800 confocal microscope.

## Results

### Single-cell RNA-Sequencing revealed seven cell types in the neurohypophysis and intermediate lobe

The pituitary is located within a bony structure of the mouse skull, dubbed *sella turcica*, allowing accurate surgical isolation of this tissue. In particular, the medially located NH can be readily observed owing to its conspicuous white color, due to the high density of neurohypophyseal axons and pituicytes. We took advantage of these anatomical features to dissect neurohypophyseal tissue from 3-month-old C57/BL6 male mice and thereafter performed scRNA-Seq analysis. Notably, the isolated tissue contained residual tissue from the adjacent intermediate pituitary lobe (IL), hence we took into consideration that our NH tissue preparation will contain some IL cells (Fig. 1A).

**Figure 1.**
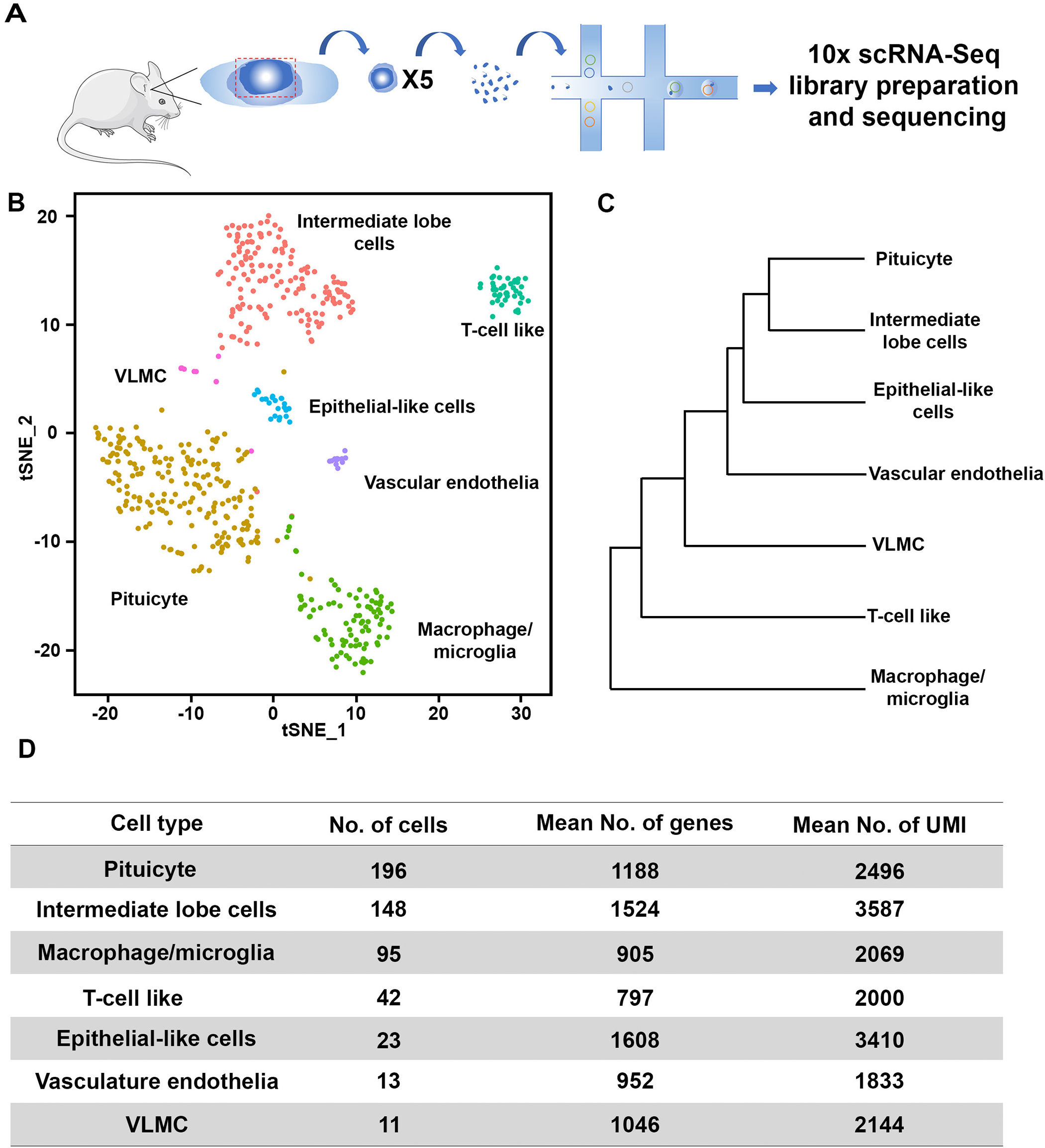
Single-cell RNA-Seq reveals seven cell types of dissected mouse NH. **(A)** Schematic representation of the scRNA-Seq procedure. Neurohypophyseal tissues were dissected from five C57BL6 adult male mice and pooled. Two independent pools were separately subjected to single cell dissociation, single cell capturing, and library preparation on 10x Chromium platform. The two libraries were then indexed and combined for sequencing using NextSeq 500 High Output v2 kit (75 cycles). **(B)** The two libraries were pooled and mapped on tSNE plot, showing cell clusters of intermediate lobe cells, T-cell like, vascular and leptomeningeal cell (VLMC), epithelial-like cells, vascular endothelia macrophage/microglia and pituicyte. Each dot represents one cell, cells with the same color belong to one cell type. **(C)** Dendrogram showing the distance matrix from the PCA space of the ‘average’ cell among the 7 cell types. The length of path between each two cell types indicates the relativeness between them. **(D)** A table summarizing the number of cells, average number of genes and UMIs found in each cell type.

We collected two pools of dissected neurohypophyseal tissue, each contained 5 mice. Single cells from the dissociated tissue were thereafter captured using the 10X Chromium gel beads in a droplet, followed by independent library preparation for each pool. The two sets of libraries were indexed and sequenced together (Fig. 1A). Low variation between the pools was detected in the PCA plot containing the two first PCs and in the tSNE plot using Seurat R package (Butler et al., 2018). The two data sets were pooled and cell clusters were built using the 900 most variable genes using FindClusters function in Seurat package using 11 principle components (PCs) and resolution 1.0 and analyzed together to create the tSNE plot (Fig. 1B and Extended Fig. 1-1). The normalized differentially expressed genes of each cluster (**Table 1-1** and **Table 1-2**) were used to identify seven major cell types, which were designated based on expression of published marker genes (Fig. 1B). Thus, we compared our gene lists to existing single-cell database, such as the mouse brain atlas from the Linnarsson Lab (Zeisel et al., 2018), the Cell type function from Allen Brain Atlas (http://celltypes.brain-map.org/), mouse vascular and vascular associated cell single-cell database (He et al., 2018; Vanlandewijck et al., 2018) and the DropViz web tool (Saunders et al., 2018). We also compared our data to published scRNA-Seq of anatomically adjacent tissues, such as the hypothalamus and the median eminence (Campbell et al., 2017; Chen et al., 2017). The identified NH cell type were labeled as: pituicyte, macrophage/microglia, vascular endothelia, T-cell like and vascular and leptomeningeal cells (VLMC). As expected, due to the nature of the dissection procedure mentioned above, we also identified IL cells. The latter was identified by comparing to recently published whole mouse pituitary single cell transcriptomes (Cheung et al., 2018; Ho et al., 2018; Mayran et al., 2018). To determine the relativeness of the clustered cell types, we used the BuildClusterTree function in Seurat R package to generated dendrogram, representing a phylogenetic tree relating the ‘average’ cell from each identity class (Fig. 1C). The number of cells, as well as mean number of genes and average number unique molecular identifiers (UMI) representing each of the designated cell types are shown in Figure 1D. Notably, the cell number does not necessarily reflect the compositional proportion in the tissue but probably randomized sampling in single cell capturing, varied resilience of different cell types to dissociation procedure and cell type-specific RNA stability.

Following the identification of NH and IL cell types we searched for sets of genetic markers characterizing each cell type. We generated a heatmap showing cluster analysis of the top twenty differentially expressed genes representing the transcriptomic profile of the various NH and IL cell types and selected three feature genes, which represent each cell type (Fig. 2). These included known markers for VLMC cells (*Ogn, Lum* and *Dcn*), fenestrated vascular endothelia (*Emcn, flt1* and *Plvap*), T-cell like (*Ms4a4b* and *Cd3d*), and marcrophage/microglia (*Ctss*, *C1qa* and *Cx3cr1*) (Kindt et al., 2007; Liu et al., 2001; Marques et al., 2016; Stan et al., 1999). In the case of three of the identified cell types, namely, epithelial-like cells, pituicytes and IL cells, the majority of the commonly used markers for these cell types were not differentially expressed between the clusters. Therefore, the unique differentially expressed featured genes we assigned for these cell types are novel markers. Thus, the novel markers *Lcn2, cyp2f2* and *Krt18* represented epithelia-like cells; *Pcks2, Scg2* and *Chga* marked IL cells, and finally, *Col25a1, Mest* and *Srebf1* were selected as pituicyte-specific markers (Fig. 2). The specificity of the selected marker genes is exemplified in Figure 3 in which a featured gene from each cluster was embedded onto the tSNE plot showing distinct distributions of different cell types (Fig. 3A). A violin plot showing the normalized log-transformed single cell expression of selected featured genes in the different cell types is shown in Figure 3B.

**Figure 2.**
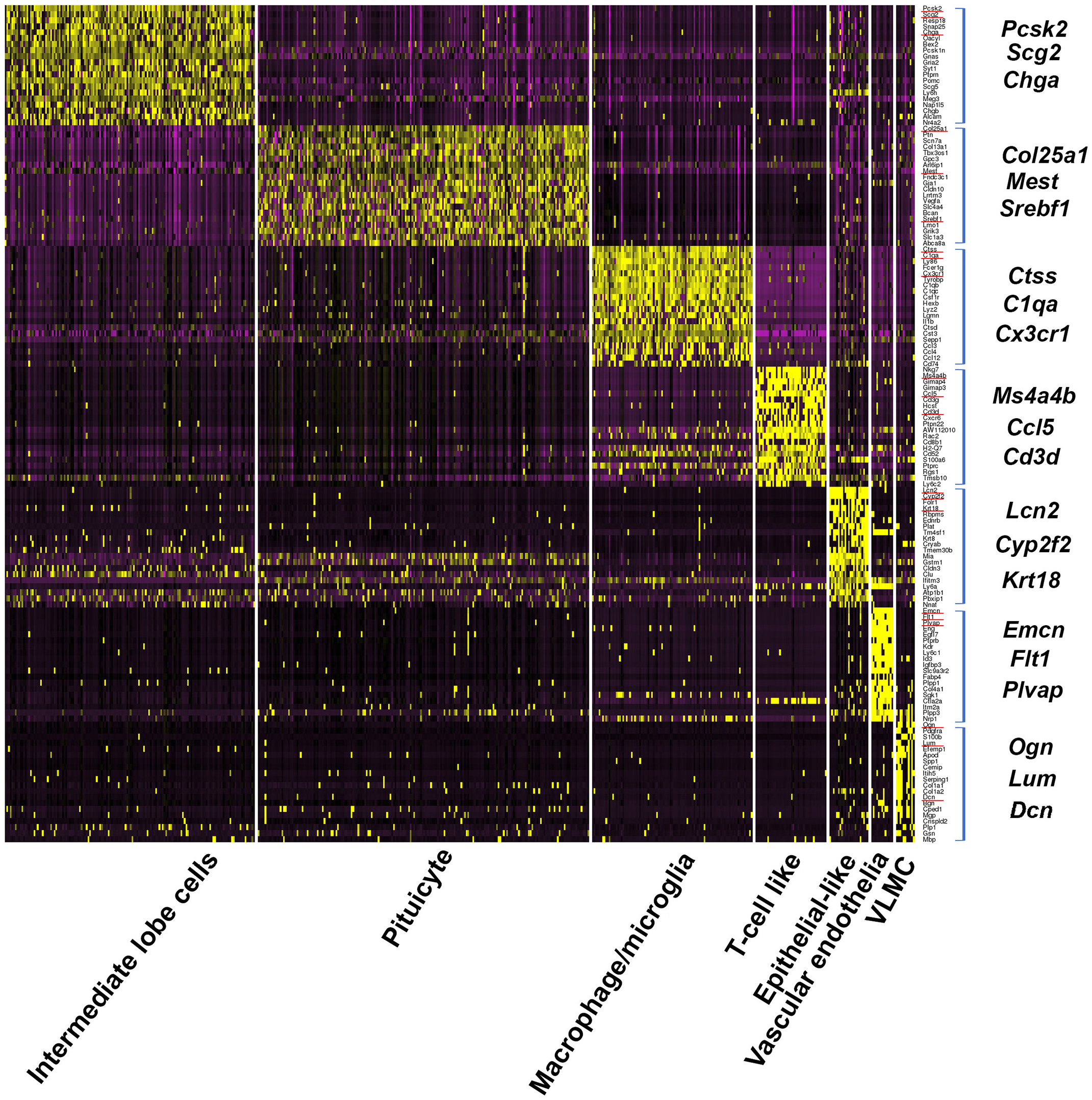
Heatmap of differentially expressed genes in neurohypophyseal and intermediate lobe cell clusters. Heatmap showing scaled gene expression of the top twenty genes (square brackets) representing each of the seven cell types found in the neurohypophysis and intermediate lobe. Each column display gene expression of an individual cell and genes are listed in the rows. Selected marker genes are underlined in red and enlarged on the side.

**Figure 3.**
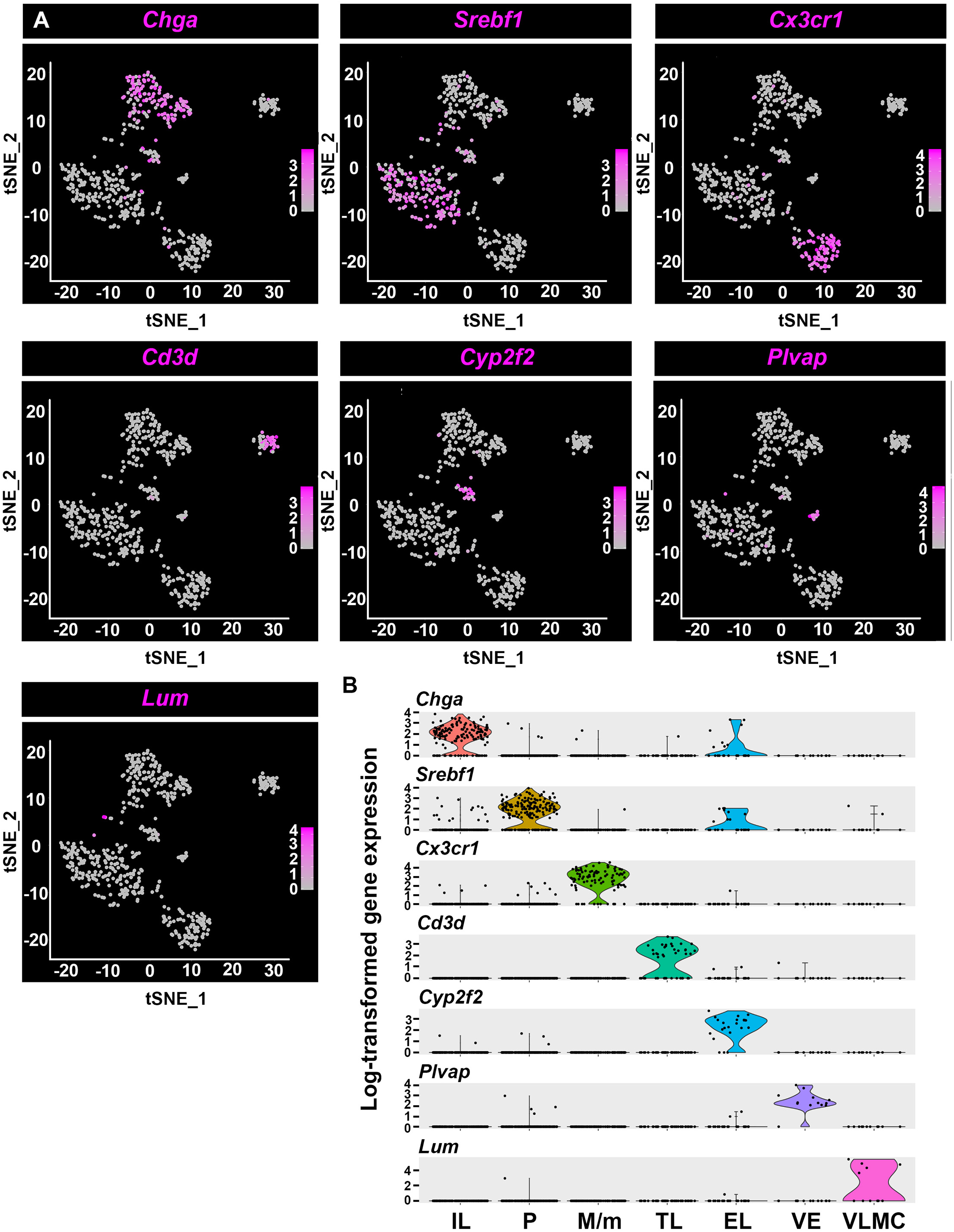
Featured genes representing the landscape of the seven neurohypophyseal and intermediate lobe cell types. **(A)** Distribution of featured genes from each cell type embedded in tSNE plots. The gene expression scale was color coded with high expression level in deep blue, low expression in gray. **(B)** Violin plots displaying normalized log-transformed expressions of each featured gene distributed across all the seven clusters. EL, epithelial-like cells; IL, intermediate lobe cells; M/m, macrophage/microglia; P, pituicyte; TL, T-cell like; VE, vascular endothelia; VLMC, vascular and leptomeningeal cells.

### Novel pituicyte genes display higher specificity than commonly used markers

We report five selected differentially expressed genes, *Srebf1, Rax, Mest, Adm* and *Col25a1*, which showed robust expression in the majority of pituicyte population (Fig. 4A). Four of these genes, *Srebf1, Rax, Adm* and *Col25a1* were robustly expressed in the pituicyte population. *Srebf1* displayed with residual expression in a small number of epithelial-like cells, but was not differentially expressed in this cluster. (Fig. 4A, **Table 1-1** and **Table 1-2**). Similarly, *Mest*, was also found to be expressed in some IL cells, but was below the differentially expression statistical significance criteria (Fig. 4A, **Table 1-1** and **Table 1-2**).

**Figure 4.**
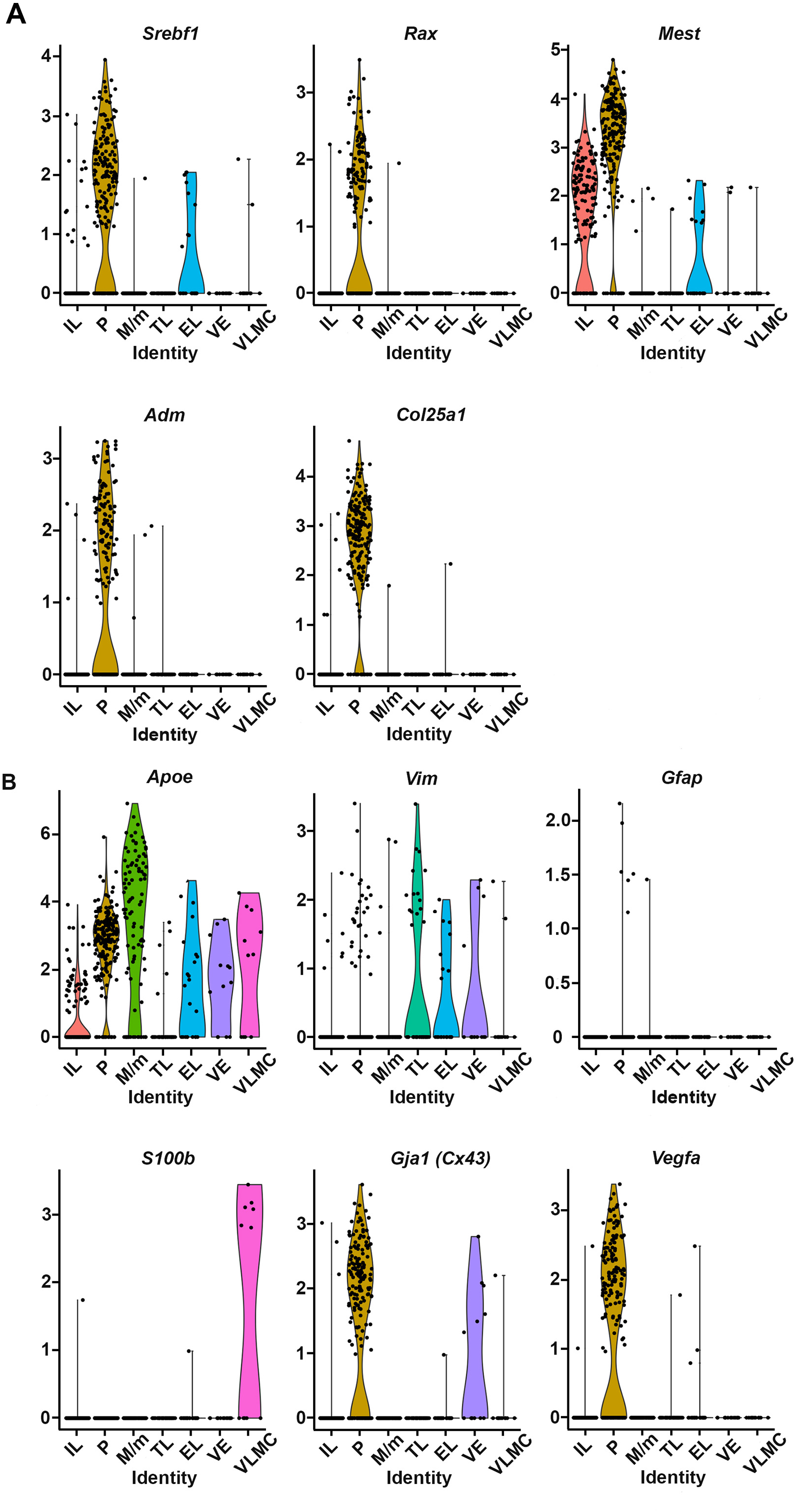
Novel pituicyte markers show higher specificity and robustness compared to previously used markers. **(A)** Violin plots displaying expression distributions of novel pituicyte marker genes in seven pituitary cell types seven clusters. *Srebf1, Rax, Mest, Adm and Col25a1* were selected from this single cell RNA-Seq data and mapped onto the violin plots. The y axis represents the normalized log-transformed expression of respective genes. Each dot represents a cell and the shape of violin represents the proportion of cells being enriched comparing to the rest of cells in a given cluster. **(B)** Previously published pituicyte markers *Apoe, Vim, Gfap, S100β*, *Gja1 (Cx43)* and *Vegfa* were mapped onto the violin plots within the seven identified cell types. EL, epithelial-like cells; IL, intermediate lobe cells; M/m, macrophage/microglia; P, pituicyte; TL, T-cell like; VE, vascular endothelia; VLMC, vascular and leptomeningeal cells.

We noticed that the novel pituicyte genes revealed by scRNA-Seq displayed higher specificity than previously published pituicytes markers (Boyles et al., 1985; Cocchia, 2004; Marin et al., 1989; Suess and Pliška, 1981; Yamamoto and Nagy, 1993). Thus, violin plots of our scRNA-Seq indicated that two commonly used pituicyte markers *Gfap* and *S100β* displayed low normalized log-transformed expressions in the pituicyte population. Furthermore, *Apoe*, which is often used as pituicyte and astrocyte marker displayed low cell-type specificity, as it was detected in all neurohypophyseal types except for T-cell like. The other three reported pituicyte markers *Gja1/Cx43*, *Vegfa* and *Vim* (Yamamoto and Nagy, 1993; Furube et al., 2014; Marin et al., 1989;) displayed higher normalized pituicyte expression and were somewhat more specific than *Apoe* (Fig. 4B). Notably, although *Vim* displayed some expression in the pituicyte cells, it didn’t pass the differentially expressed criteria in pituicyte cluster comparing to other cell types (**Table 1-1** and **Table 1-2**).

We next examined whether the novel pituicyte markers identified by scRNA-Seq are expressed in the mouse NH by *in situ* hybridization. The selected pituicyte marker *Col25a1* with robust normalized expression (adjusted p-value 8.38E-82, average log_2_ fold change 1.67) in the pituicyte cluster according our scRNA-Seq analysis was subjected to wholemount mRNA *in situ* hybridization, followed by immunostaining with an antibody against the previously published pituicyte marker *Vim* (Fig. 4A and **Table 1-2**). This analysis showed that *Vim* immunoreactivity is detected in a subset of *Col25a1*-positive cells (Fig. 5 and **Extended Video 5-1**). This analysis was in agreement with our scRNA-Seq bioinformatic analysis (Fig. 4), suggesting that some of the commonly used pituicyte markers also label other NH cell types.

**Figure 5.**
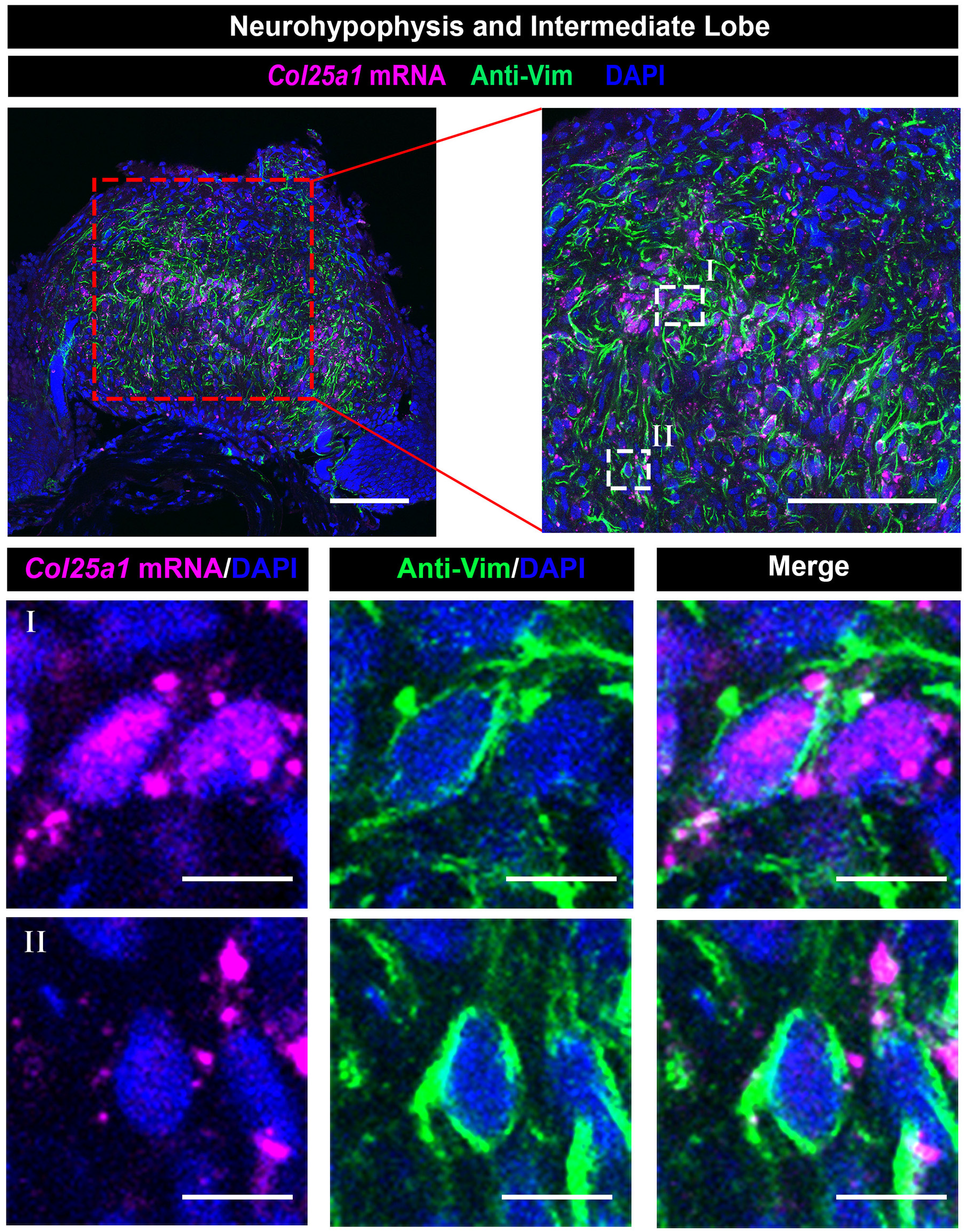
Expression of the novel pituicyte marker *Col25a1*, in the neurohypophysis. Validation of the scRNA-Seq results using wholemount staining of dissected neurohypophysis (NH) derived from a C57/BL6 adult mouse. Dissected NH was subjected to fluorescent mRNA *in situ* hybridization with an antisense *Col25a1* probe, followed by immunostaining with an antibody directed to the Vim protein and visualized by confocal microscopy. The top panels display different magnifications (scale bars = 100µm) a single confocal optical plane of *Col25a1, Vim* and the nuclei dye, DAPI. Highly magnified field (scale bars = 10µm) of views showing a representative *Col25a1****^+^****; Vim****^+^*** pituicyte (I) and another *Col25a1****^-^****; Vim****^+^*** neurohypophyseal cell (II).

### Spatial organization of neurohypophyseal cell types

To better understand the spatial organization of neurohypophyseal cell types, we analyzed the expression of selected genetic markers representing the major NH cell types we identified and localized the expression on horizontal section of whole mouse pituitary (Fig. 6A). We performed single molecule fluorescent *in situ* hybridization (smFISH) on a transgenic macrophage/microglia reporter (Fig. 6) as well as wholemount mRNA *in situ* hybridization combined with antibody staining (Fig. 7). Our scRNA-Seq analysis indicated that *Srebf1* is a novel pituicyte marker displaying limited expression in the epithelial-like cells, while *Cyp2f2* was exclusively expressed in epithelial-like cells (Figs. 2-4). Accordingly, *Srebf1* was specifically expressed in the NH (Fig. 6B I), while *Cyp2f2-*expressing cells were mostly located at the boundary between the IL and the AH (Fig. 6B II). Moreover, *Cyp2f2-*positive cells were not found in the NH suggesting that the epithelial-like cells we identified using scRNA-Seq were likely derived from the IL and not from the NH (Fig. 6B II&III and Extended Fig. 6-1). This conclusion was further confirmed using smFISH to probe another specific epithelial-like featured gene, *Lcn2*, which was specifically expressed by IL cells but not in the NH (Extended Fig. 6-2A)

**Figure 6.**
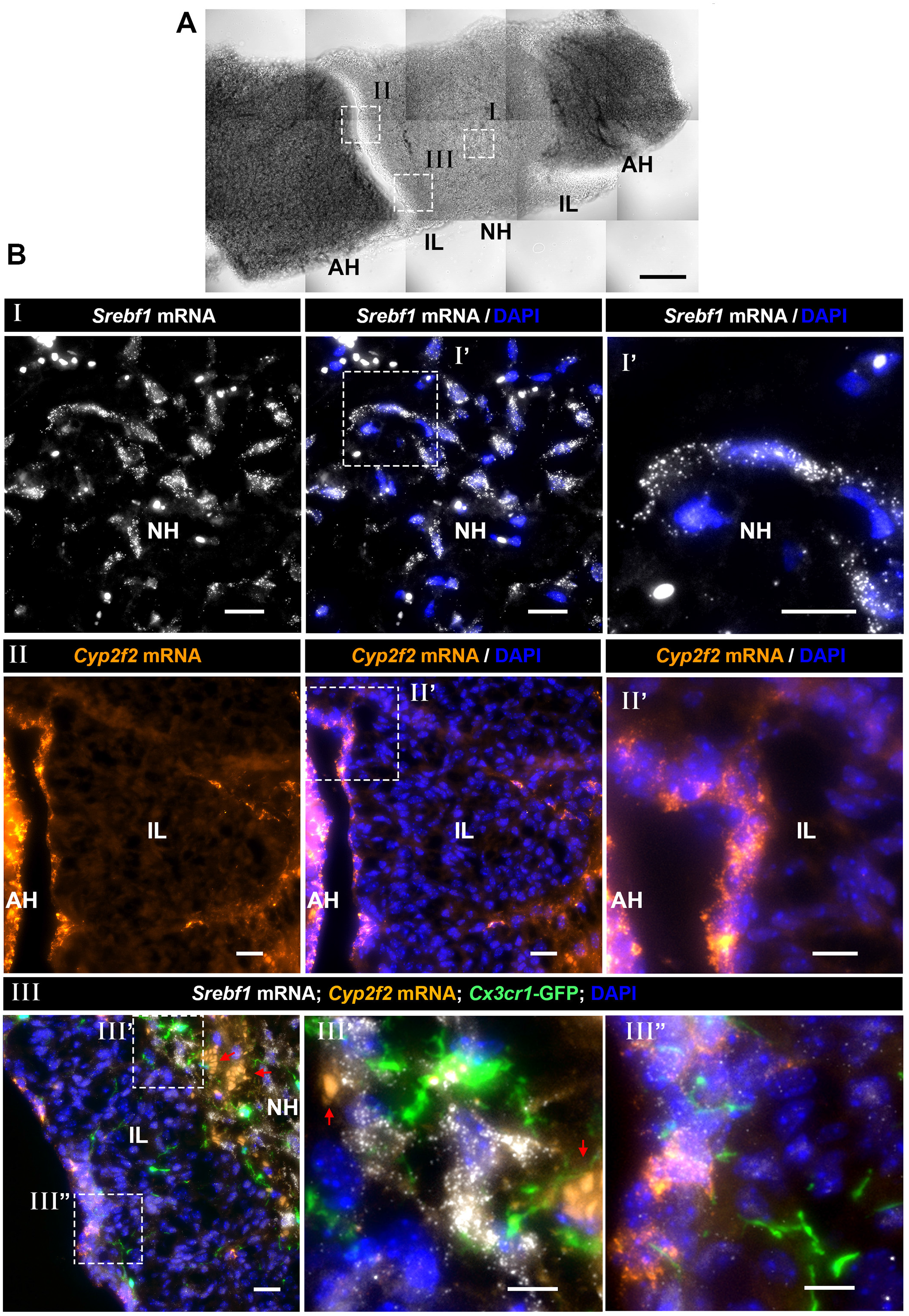
Spatial distribution of pituicyte, macrophage/microglia and epithelial-like cells in the neurohypophysis and intermediate lobe. **(A)** A brightfield image of a horizontal section of adult mouse pituitary showing the locations of the neurohypophysis (NH), intermediate lobe (IL) and adenohypophysis (AH). The white boxes in the brightfield image mark the locations of specific pituitary sub-domains shown in the fluorescent images below. Scale bar, 100µm. **(B)** Different fields of views (marked by roman numbers) of horizontal section (7µm) of pituitaries derived from three-month-old Cx3cr1-GFP macrophage/microglia transgenic reporter mouse, which were subjected to single molecule fluorescent *in situ* hybridization (smFISH) with antisense probes directed to *Srebf1* (I), *Cyp2f2* (II) or multiplexed smFISH of *Srebf1* and *Cyp2f2* on *Cx3cr1*:GFP mouse (III) to observe the relative location of selected cell types. Scale bars equal to 20µm. High magnification image of the region delineated with white dashed box are shown. Scale bar equals to 10µm. (II). Note that the smFISH probe of epithelial-like cell marker, *Cyp2f2*, labels the border between the IL and the AH, as well as IL cells. Arrows indicate background autofluorescent signals of circulating erythrocytes. Scale bars, 20µm in Panels (I) and (II); 10µm in panel (III).

**Figure 7.**
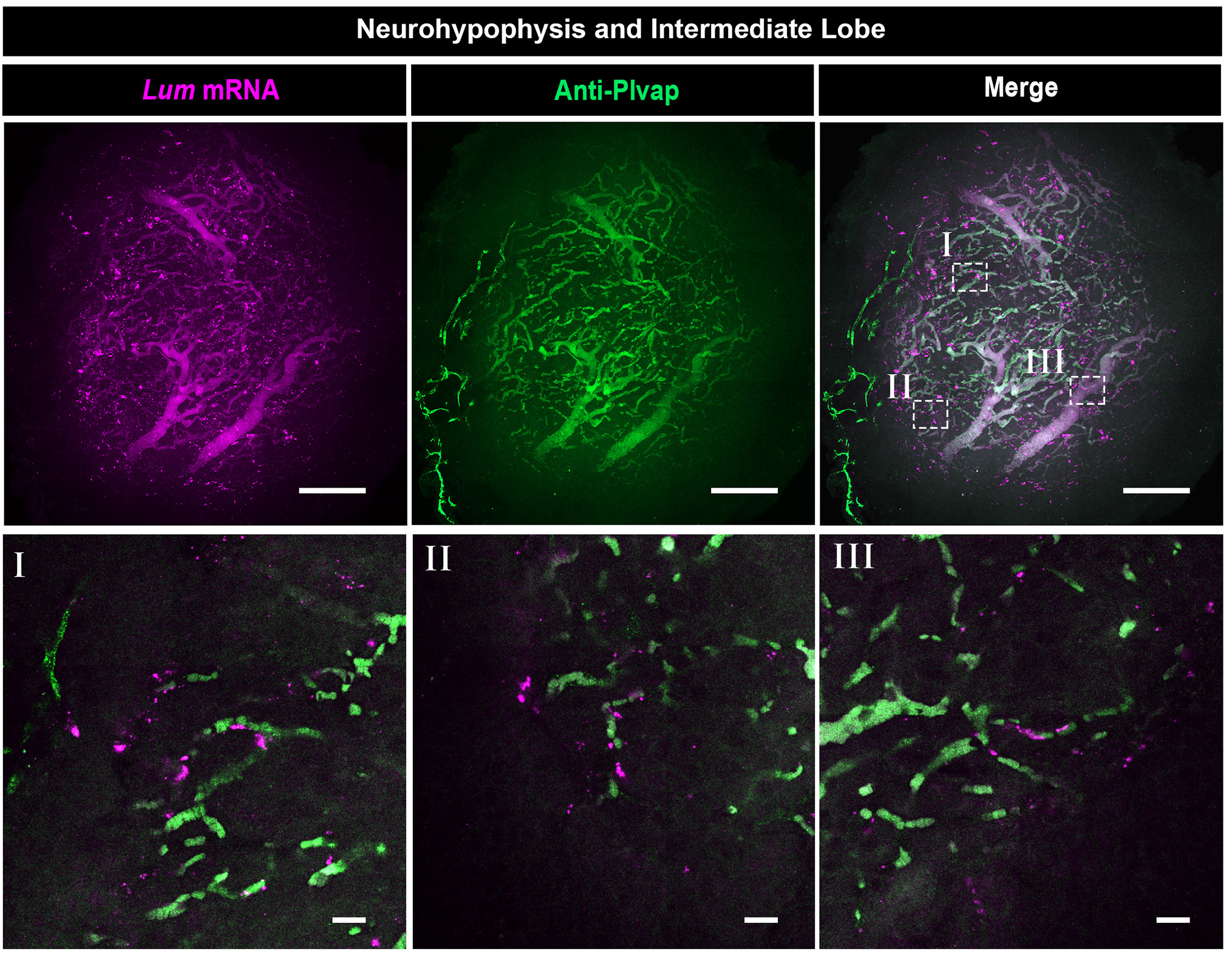
Neurohypophyseal VLMCs are associated with fenestrated vascular endothelia. Confocal Z-stack (maximum intensity projection) of dissected neurohypophysis, which was to wholemount fluorescent *in situ* hybridization with an antisense RNA probe directed to the VLMC marker, *Lum*, followed by immunostaining with an antibody directed to *Plvap* protein, which is a fenestrated endothelia marker (Scale bars = 100µm). The bottom panels (labeled I-III) display high-magnification single plane confocal images of the of respective regions delineated in white boxes in the top right panel (Scale bars = 20µm).

We next performed simultaneous labeling of macrophage/microglia, pituicyte and epithelial-like cells by performing smFISH of *Srebf1* and *Cyp2f2* probes on pituitaries of transgenic *Cx3cr1*:GFP reporter mice, labelling macrophage/microglia (Jung et al., 2000). We observed that the *Cx3cr1*: GFP-positive macrophage/microglia were distributed throughout the whole pituitary, including the NH, IL and AH (Fig. 6B III and Extended Fig. 6-2B). These macrophages/microglia were intermingled with both *Srebf1****^+^****; Cyp2f2****^-^*** pituicytes and *Cyp2f2****^+^*** epithelial-like cells suggesting possible functional cell-cell interactions between pituitary cells and these macrophages/microglia (Fig. 6B III’ & III”). Some of these *Cyp2f2^+^* IL cells co-expressed *Srebf1*, however they were spatially separated from pituicytes located in the NH (Fig. 6B III and Extended Fig. 6-1). This result is in agreement with our bioinformatic scRNA-Seq analysis showing that a small number of epithelial-like cells expressed *Srebf1* (Fig. 4A).

Our scRNA-Seq analysis also detected a NH cell population, which co-expressed *Pdgfra* and *Lum* (Fig. 2 and **Table 1-2**). We assumed that this cell population is similar or identical to the so called vascular and leptomeningeal cell (VLMC), which has been found to localized on blood vessels of the brain (He et al., 2018; Marques et al., 2016; Vanlandewijck et al., 2018). We therefore examined the tissue distribution of VLMC cells and fenestrated neurohypophyseal vascular endothelia, which express the *Plvap* protein (Stan et al., 1999). This analysis confirmed that as in the case of brain vasculature, VLMCs were in close association with the fenestrated endothelia of the NH (Fig. 7 I-III). Finally, although we have found a small population of T-cell like cells in the NH, immuno-staining of the T-cell-specific cell surface marker, *Cd3*, revealed low abundance of *Cd3-*poistive cells in the NH (Kindt et al., 2007) (Extended Fig. 6-2C). It is likely that these T cells are not resident NH population but rather a transient population, which is transported from the blood. The above *in situ* hybridization analyses confirmed our featured gene designation determined by scRNA-Seq.

Taken together, Our gene expression analysis of NH and IL reveal a comprehensive view of neuro-intermediate lobe cell types in adult male mice. This study provides an important resource for specific functional studies and cell-cell interactions between the various NH cell types.

## Discussion

The NH is a major neuroendocrine interface, which allows the brain to regulate the function of peripheral organs in response to specific physiological demands. Despite its importance, a comprehensive molecular description of cell identities in the NH is still lacking. Recent studies revealed cell heterogeneity of whole pituitary gland using scRNA-Seq, however, these studies did not separated the NH from the adjacent adenohypophysis and very few NH cells with limited sequence information were reported (Cheung et al., 2018; Ho et al., 2018; Mayran et al., 2018). Here, utilizing scRNA-Seq technology, we identified the transcriptomes five major neurohypophyseal cell types and two IL cell populations in the adult male mice. Using selected featured genetic markers, we mapped the spatial distribution of selected cell type *in situ*.

The identified differentially expressed gene clusters revealed by scRNA-Seq correspond to previously characterized cell types. Thus, previous studies reported the appearance of pituicyte (Anbalagan et al., 2018; Furube et al., 2014; Seyama et al., 1980; Wittkowski, 1986; Yamamoto and Nagy, 1993), macrophage/microglia (Kindt et al., 2007; Pow et al., 1989; Sasaki and Nakazato, 1992), and fenestrated endothelia (Stan et al., 1999) in the NH. The identification of IL and epithelial-like cells in the present study is in agreement with other reports of these cells in the mammalian pituitary (Hudson, 2002; Moran et al., 2010) and also matches recent scRNA-Seq analyses of whole mouse pituitary (Cheung et al., 2018; Ho et al., 2018; Mayran et al., 2018).

The novel pituicyte markers identified in our study showed much more specific and robust expression than all previously published pituicyte markers. Among them, *Mest, Srebf1, Adm* and *Col25a1* were not reported to be expressed by pituicyte before. *Vegfa* was reported as pituicyte marker in both mice and zebrafish (Anbalagan et al., 2018; Furube et al., 2014). Another prominent pituicyte marker we identified, namely *Rax*, is a known hypothalamic tanycyte marker which is also expressed in the NH (Miranda-Angulo et al., 2014; Pak et al., 2014). We show here that *Rax* is expressed in pituicyte which is in line with the notion that tanycytes and pituicytes are of common astrocytic lineage (Clasadonte and Prevot, 2018). *Adm*, another prominent pituicyte marker, which has been attributed to rat hypothalamic paraventricular nuclei and supraoptic nuclei neurons rather than glia (Ueta et al., 1999). Finally, in agreement with our findings, *Col25a1* was found to be enriched in neurohypophysis via the database Bgee (Bastian et al., 2008).

Our novel pituicyte markers displayed greater specificity (i.e. adjusted p-value < 0.05 for differential expression), higher expression level (average log_2_ fold change > 1) and robustness (i.e. abundance in pituicytes) compared to the most commonly used markers. Thus, as we previously showed in the case of zebrafish pituicytes (Anbalagan et al., 2018), we found that *Apoe* is broadly expressed in multiple mouse NH cell types. However, although *Vim* and *Gfap* displayed relatively low mRNA expression level in our scRNA-Seq analysis, their protein immunoreactivity was readily detectable in the NH. This could be due to the inherently shallow sequencing method for 10x Genomics platform. The astroglial protein *S100β* is also used to label pituicytes (Cocchia, 2004). It was reported that *S100β* is highly abundant when compared to *Vim^+^* and *Gfap^+^* cells (Virard et al., 2008; Wei et al., 2009). However, in our study, *S100β* was not among the top differentially-expressed pituicyte genes, but was found to be exclusively expressed in VLMC cell type. In view of the gene coverage limitation of the 10x Genomics platform, *S100β* might have been missed in our analysis, hence, future studies should be aware of our finding regarding its expression in VLMC. Another known pituicyte-specific marker, namely *Gja1*, also known as *Cx43*, displayed robust specific expression in our mouse pituicyte cluster. This is in agreement with the reported findings in rat and zebrafish (Anbalagan et al., 2018; Yamamoto and Nagy, 1993).

We identified VLMC as a new neurohypophyseal cell type which is marked by the prominent expression of *Pdgfra* and *Lum.* We further showed that VLMC are associated with *Plvap^+^* fenestrated neurohypophyseal capillaries. In agreement with our findings, *Pdgfra^+^*;*Lum*^+^ VLMC population was found in the mouse brain as vascular-associated cell type (Marques et al., 2016) or as fibroblast-like cells that are loosely attached to vessels and located in between smooth muscle cells and astrocyte end-feet (Vanlandewijck et al., 2018). Although VLMC express some markers of oligodendrocyte precursor cells (OPCs), such as *Pdgfra*, they are distinct from OPCs and oligodendrocyte lineages (Marques et al., 2016).

Importantly, previous reports have described the existence of OPCs in the NH (Miyata, 2017; Virard et al., 2006, 2008). We did not detect OPCs in the present study, however, this could be due to low abundance of these cells in our tissue. Alternatively, because Virard et al. relied on *Pdgfra* as a sole OPC marker, it is possible that they misidentified VLMCs as OPCs. Notably, Virard et al. reported that these *Pdgfra*^+^ cells were shown to be pituicyte progenitors in their study (Virard et al., 2006). Similarly, other studies reported that VLMC display multipotent stem cell niche function in the CNS and other organs suggesting that they may play a similar roles in NH function (Nakagomi and Matsuyama, 2017; Ueharu et al., 2018). Further studies are required to determine whether neurohypophyseal OPCs are in fact VLMCs and if VLMCs are pituicyte progenitors.

Although pericytes have been previously reported to be associated with neurohypophyseal capillaries (Miyata, 2017; Nishikawa et al., 2017), we did not detect them in the present study, possibly due to the fact that isolating pericytes requires different tissue dissociation conditions.

Macrophage/microglia were found in our study as prominent NH resident cells. Previous report showed that neurohypophyseal microglia in rat endocytose and digest axonal terminus (Pow et al., 1989), whereas the pituicyte envelops the buttons of axons (Morris, 1976) and provide cues for the permeable endothelia fate (Anbalagan et al., 2018). Our findings that macrophage/microglia are intermingled with the pituicyte in the NH are agreement with such functional cooperation between these two cell types.

In summary, our transcriptome analysis of individual cells derived from NH and IL tissues of adult male mice have revealed the cellular heterogenicity of the NH and provide novel molecular markers for the major cells in those tissues. We present a valuable resource which will serve as the basis for further functional studies.

## Acknowledgements

We thank Yael Kuperman for her help in neurohypophysis dissection; Hagit Dafni for providing C57BL6 mice for dissociation protocol optimization; Stefan Jung for providing he *Cx3cr1*:GFP mice and Shalev Itzkovitz for the smFISH probes; Amrutha Swaminathan and Ludmila Gordon for their valuable comments on the text and figures and Hanjie Li for his advices on cluster annotation. G.L. is supported by the Israel Science Foundation (#1511/16); Minerva-Weizmann program, Adelis Metabolic Research Fund, Yeda-Sela Center for Basic Research (in the frame of the Weizmann Institute), the Nella and Leon Benoziyo Center for Neurological Diseases, Richard F. Goodman Yale/Weizmann Exchange Program and Estate of Emile Mimran. G.L. is an incumbent of the Elias Sourasky Professorial Chair.

## Authors’ contributions

Q.C., and G.L. planned the project. Q.C. performed all of the experiments, analyzed and prepared the figures. D.L. and Q.C. analyzed the scRNA-Seq data. J.B. contributed to the preparation of neurohypophyseal tissues and to the following 10X Genomics Chromium scRNAseq procedure together with Q.C. Q.C. and G.L. wrote the manuscript. All authors reviewed the manuscript’s text.

---

**Extended figure 1-1.**
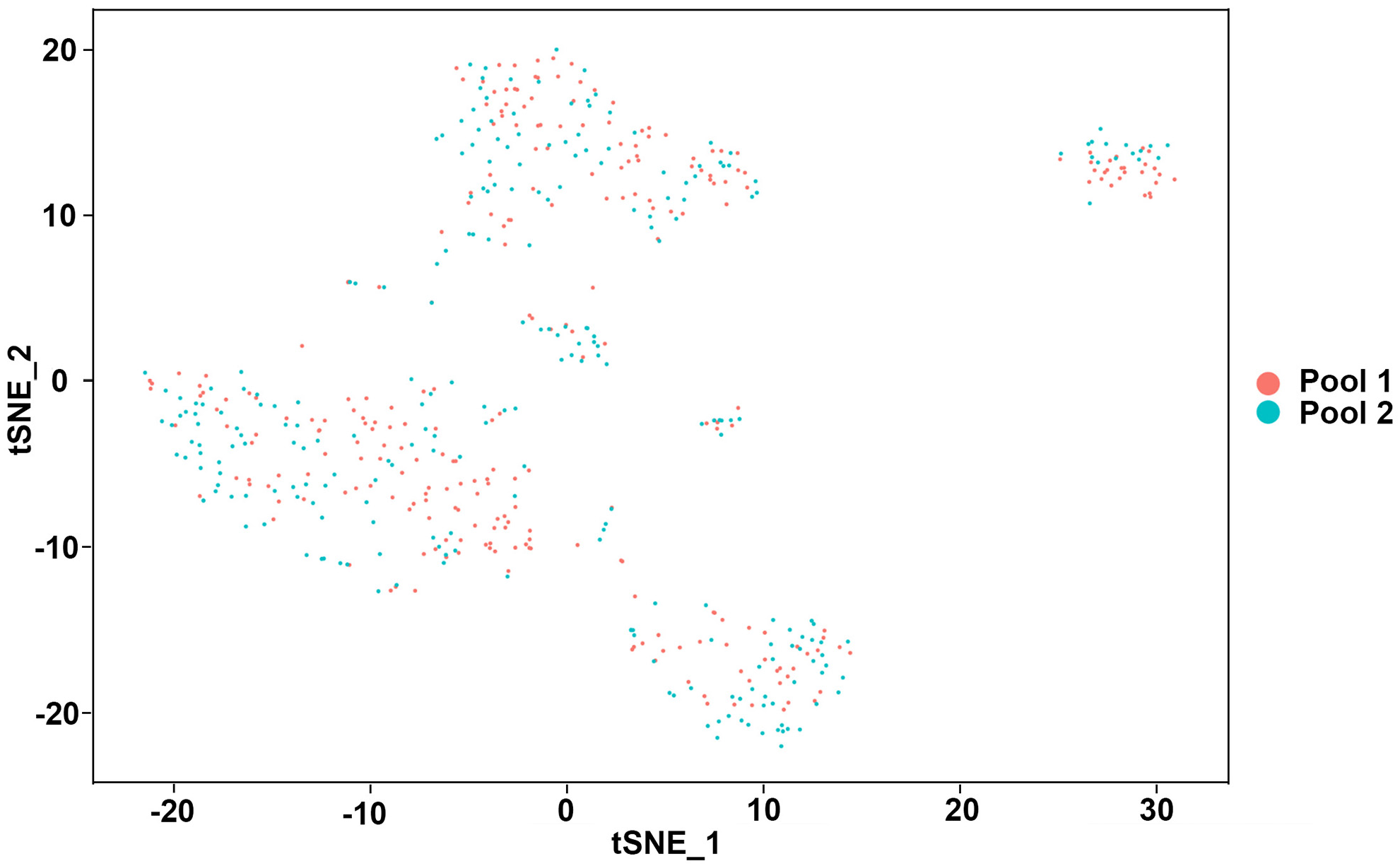
A tSNE plot shows two pools of scRNA-Seq libraries. A tSNE plot in which the cell is colored according to pool origin.

**Extended figure 6-1.**
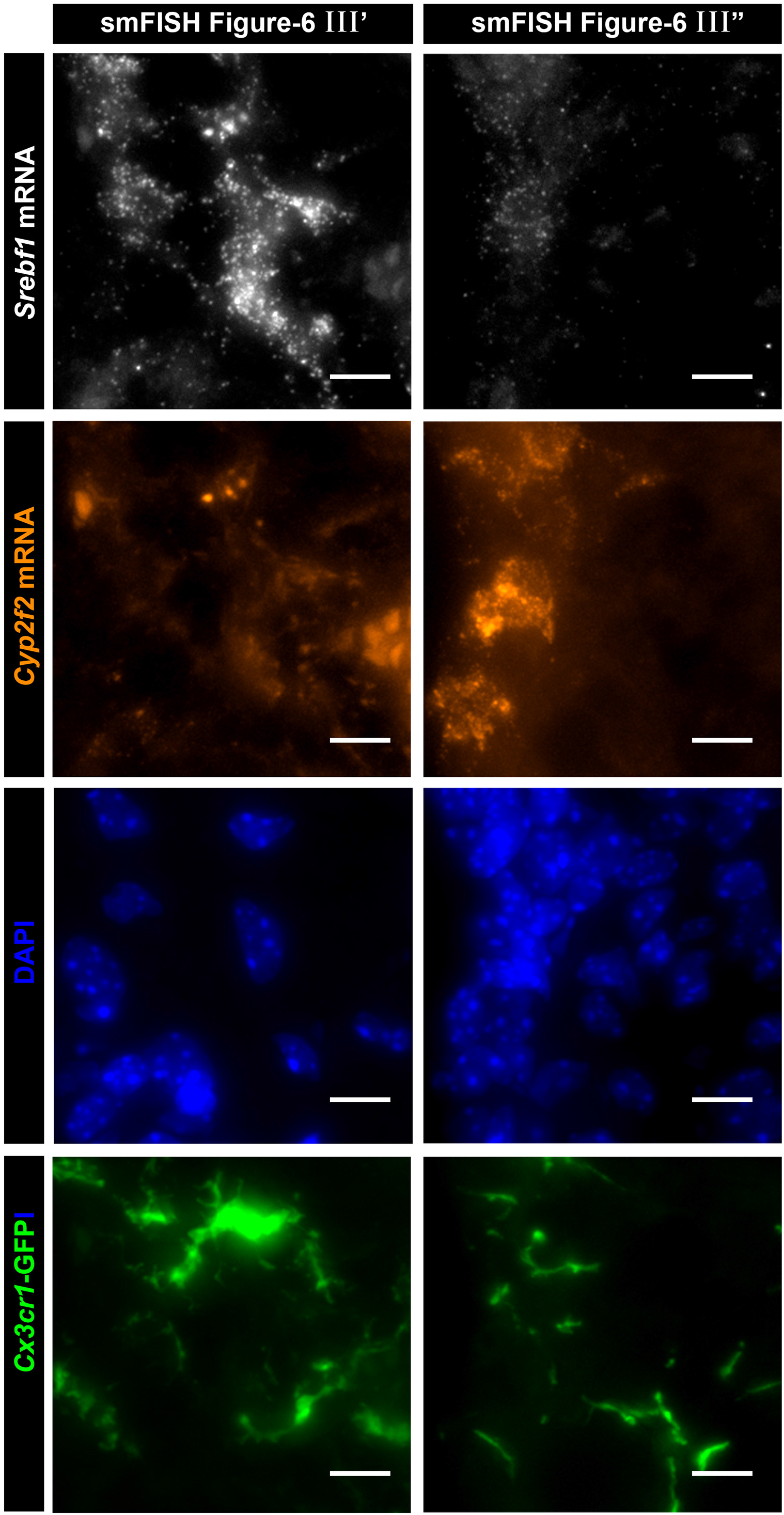
Separate channels for multiplex smFISH images in Figure 6 III’ and III”. High magnification of the images in Figure 6B III’ and III” showing horizontal section (7 µm) of pituitaries derived from a three-month-old *Cx3cr1*-GFP macrophage/microglia transgenic reporter mouse, which were subjected to single molecule fluorescent *in situ* hybridization (smFISH) with antisense probes directed to *Srebf1* and *Cyp2f2*. The purpose of this figure is to better visualize the overlapping signals of four channels representing pituicyte marker *Srebf1* mRNA, epithelial-like cells marker *Cyp2f2* mRNA, nucleus stain DAPI and macrophage/microglia (*Cx3cr1*-GFP). Scale bars, 10µm.

**Extended figure 6-2.**
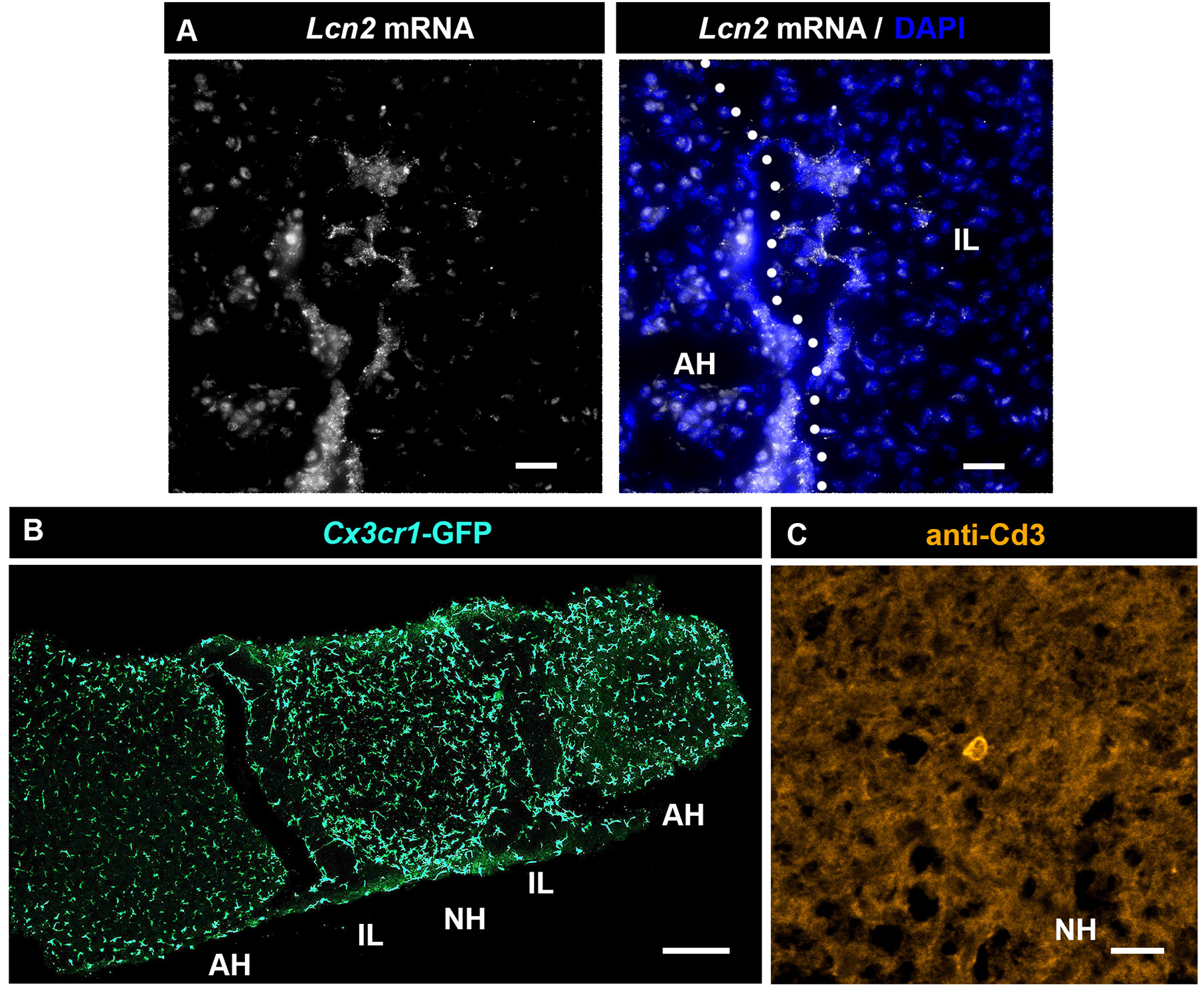
Validation of macrophage/microglia, intermediate lobe cells and T-cell like clusters. **(A)**. Location of the epithelial-like cells was further validated by smFISH using another novel marker *Lcn2* selected from the scRNA-Seq data. They were located at the boundary between the IL and the AH, which is marked by the white dotted line. Scale bars, 20µm. **(B)**. Confocal Z-stack (maximum intensity projection) of horizontal vibratome pituitary section (50µm) from a three-month-old male mouse harboring the transgenic macrophage/microglia reporter *Cx3cr1*-GFP. The macrophage/microglia located all over the pituitary. Scale bar, 200µm. **(C)**. Immunostaining of the T-cell marker, *Cd3*, showing a lone T-cell inside the NH horizontal pituitary cryosection (7µm) of a 3-month-old C57/BL6 male mouse. Scale bar, 20µm.

**Table 1-1. Complete list of normalized differentially expressed genes**

A list of normalized differentially expressed genes that were expressed in at least 25% of the cells within between the two groups of cells.

**Table 1-2. Filtered list of normalized differentially expressed genes**

List of normalized differentially expressed genes in each cell type displaying an average log_2_ fold change > 1 and adjusted *p*-value < 0.05.

**Extended video 5-1.**

Whole mount *in situ* hybridization of *col25a1* co-stained with Vim antibody and DAPI on dissected neurohypophysis of 3-month-old C57/BL6.

## Notes

**Conflict of interest:** the authors declare no competing financial interests.

